# Vascular proteome responses precede organ dysfunction in sepsis

**DOI:** 10.1101/2021.12.07.471579

**Authors:** James T. Sorrentino, Gregory J. Golden, Claire Morris, Chelsea Painter, Victor Nizet, Alexandre Rosa Campos, Jeffrey W. Smith, Christofer Karlsson, Johan Malmström, Nathan E. Lewis, Jeffrey D. Esko, Alejandro Gómez Toledo

**Affiliations:** Bioinformatics and Systems Biology Graduate Program, University of California, San Diego, La Jolla, CA, USA; Department of Bioengineering, University of California, San Diego, La Jolla, CA, USA; Department of Cellular and Molecular Medicine, University of California, San Diego, La Jolla, CA, USA; Glycobiology Research and Training Center, University of California, San Diego, La Jolla, CA, USA; Department of Pediatrics, University of California, San Diego, La Jolla, CA, USA; Skaggs School of Pharmacy and Pharmaceutical Sciences, University of California, San Diego, La Jolla, CA, USA; The Cancer Center and The Inflammatory and Infectious Disease Center, Sanford-Burnham-Prebys Medical Discovery Institute, La Jolla, CA, USA; Department of Clinical Sciences, Division of Infection Medicine, Lund University, BMC, Lund, Sweden; National Biologics Facility, Technical University of Denmark, Krogens-Lyngby, Denmark

**Keywords:** Vascular glycocalyx, proteome, sepsis, *Staphylococcus aureus*

## Abstract

Vascular dysfunction and organ failure are two distinct, albeit highly interconnected clinical outcomes linked to morbidity and mortality in human sepsis. The mechanisms driving vascular and parenchymal damage are dynamic and display significant molecular crosstalk between organs and tissues. Therefore, assessing their individual contribution to disease progression is technically challenging. Here, we hypothesize that dysregulated vascular responses predispose the organism to organ failure. To address this hypothesis, we have evaluated four major organs in a murine model of *S. aureus* sepsis by combining in vivo labeling of the endothelial proteome, data-independent acquisition (DIA) mass spectrometry, and an integrative computational pipeline. The data reveal, with unprecedented depth and throughput, that a septic insult evokes organ-specific proteome responses that are highly compartmentalized, synchronously coordinated, and significantly correlated with the progression of the disease. Vascular proteome changes were found to precede bacterial invasion and leukocyte infiltration into the organs, as well as to precede changes in various well-established cellular and biochemical correlates of systemic coagulopathy and tissue dysfunction. Importantly, our data suggests a potential role for the vascular proteome as a determinant of the susceptibility of the organs to undergo failure during sepsis.

## Introduction

Sepsis is a dysregulated host response to infections that can rapidly evolve into a life-threatening pattern of multiple organ dysfunction(1). The disease burden of sepsis has been estimated to ∼50 million incident cases and ∼11 million fatalities each year, which accounts for almost 25% of all annually reported global deaths(2). Despite these shockingly high levels of morbidity and mortality, no specific sepsis biomarkers are currently available. Treatments remain generic and limited to broad-spectrum antibiotics therapy, intravenous fluid resuscitation, and supportive care(3, 4). The molecular basis of disease progression is also poorly understood, hindering the early recognition and management of critically ill patients in the intensive care units (ICUs).

Vascular dysfunction and organ failure are two highly intertwined clinical outcomes that determine most of the morbidity and mortality of human sepsis(5). Being a systemic syndrome, sepsis is characterized by extensive molecular crosstalk between dysregulated inflammatory, coagulopathic and metabolic processes, operating both at the local and systems levels, and compartmentalized across the blood-tissue interfaces of the organs. Therefore, dissecting the individual contribution of vascular dysfunction to the development of organ damage is a difficult task. Nevertheless, understanding the spatiotemporal relationships between vascular and parenchymal tissues at the molecular level might unveil novel ways to impact sepsis outcomes. For example, pharmacological stabilization of the angiopoietin (Ang)/Tie2 axis reduces vasculopathy and inflammation in preclinical models of sepsis, directly implicating endothelial dysfunction in the etiology of organ failure and emphasizing the vasculature as a legitimate therapeutic target(6–8). However, the temporal and causal links between vascular dysfunction and organ damage are still poorly understood, and new methodological frameworks are needed to capture disease progression in a compartmentalized and time-resolved fashion.

Sepsis induces dramatic perturbations in the plasma and organ proteomes of both humans and mice, highlighting the potential of proteomics as sensitive molecular readouts of disease progression(9–14). However, most of the proteomics studies to date are limited to the analysis of single time points and/or single tissue compartments. In fact, despite the recurrent observation that vascular failure lies at the core of sepsis pathology, methodological limitations still preclude the detailed study of vascular proteome changes in sepsis, and their contribution to organ damage. Vascular proteome profiling in vivo is particularly challenging, and the suitability of in vitro cellular systems has been questioned in multiple studies(15, 16). We have previously developed chemical-biology methods to tag vascular proteins in vivo, and reported that *S. aureus* sepsis triggers substantial changes in the murine vascular proteome(17, 18). However, the temporal dynamics of the vascular proteome during infection, its molecular relationships with plasma and extravascular compartments, as well as its contribution to overall sepsis progression remain unaddressed.

In this study, we hypothesize that dysregulated vascular proteome responses predispose the organism to organ failure. Using chemical-proteomics and an integrative computational pipeline, we generated a large-scale molecular description of the proteome trajectory of sepsis, in a compartmentalized and time-resolved fashion. We provide evidence for the presence of organotypic proteome changes triggered by *S. aureus* sepsis across the blood-tissue interfaces of four major murine organs (SI appendix, Fig. S1). These proteome responses are largely compartmentalized, temporally coordinated, and significantly correlated with the progression of the disease. Vascular proteome alterations were also found to precede changes in well-established cellular and biochemical correlates of sepsis, opening a window for the discovery of new diagnostic markers and novel therapeutic targets. More importantly, our data suggests a potential role for the vascular proteome as a determinant of the susceptibility of the organs to develop coagulopathy and tissue dysfunction.

## Results

### A two-staged model of the plasma proteome response to sepsis

To start tracking the progression of the host response to *S. aureus* bacteremia, mice were intravenously challenged with a LD_100_ bacterial dose, and their plasma proteomes were analyzed at 0h, 6h, 12h and 24h post infection. To facilitate accurate time-resolved quantitative proteomics, a data-independent acquisition (DIA) workflow based on Sequential Window Acquisition of all Theoretical Mass Spectra (SWATH) was applied (19). DIA-SWATH-MS uses broad isolation windows to measure almost all MS detectable peptides in a biological sample. This results in an increased number of protein identifications and more robust quantification compared to approaches based on data-dependent acquisition (DDA). Initially, 1μL of non-depleted unfractionated plasma from *S. aureus* infected or healthy control mice was subjected to proteomics analysis. Protein intensities were extracted from the DIA data using predicted spectral libraries generated through deep neural networks in the software suite DIA-NN(20), and further constrained by experimental information from public databases (Material and Methods). As shown in Fig 1A, the plasma proteome is significantly altered during infection, and clustering analysis segregates the samples into early and late responses (Fig. 1B). To uncover the main protein changes driving the partitioning of the samples, the data were deconvolved through non-negative matrix factorization (NMF)(21). Three NMF clusters or “protein signatures” were successfully identified and condensed into discrete proteome trajectories (i.e., temporal intensity patterns) (Fig. 1C). The NMF Signature 1 identified the early (6h) accumulation of inflammatory proteins, complement, coagulation factors and acute-phase reactants (APRs), including the C-reactive protein (CRP), the serum amyloid protein 2 (SAA2), and soluble CD14, as well as antimicrobial factors from the cathelicidins family (Fig. 1D). The NMF Signature 2 was characterized by the plasma accumulation of liver metabolic enzymes (Fig. 1E), whereas the NMF Signature 3 identified the depletion of hepatic factors involved in lipid metabolism (Fig. 1F). These two last signatures were linked to the late time points (12h and 24h), and were indicative of the occurrence of acute liver failure. Based on these data-driven patterns, the plasma proteome analysis reflected a two-staged model of the progression of sepsis, each stage associated with specific inflammatory and metabolic alterations. (Fig. 1G).

**Fig. 1.**
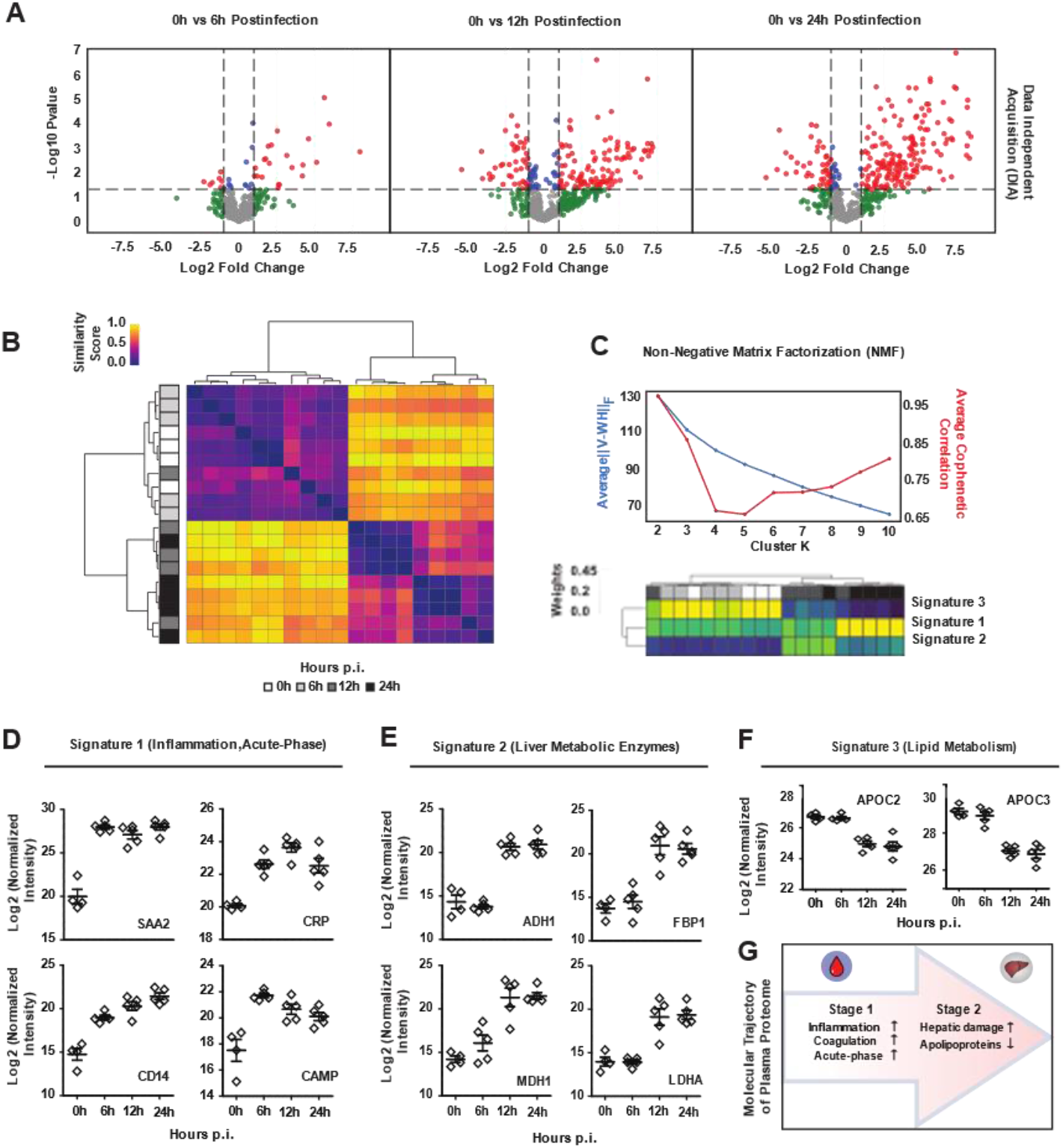
Time-resolved analysis of plasma proteome trajectories in a murine model of Staphylococcus aureus sepsis. *(A)* Volcano plots showing time-dependent changes in protein abundances in murine plasma at 0h (n=5), 6h (n=5), 12h (n=5) and 24h (n=5) post infection and captured by proteomics analysis using data-independent acquisition (DIA) mass spectrometry. Coloring thresholds were set at fold-change > 2 and P < 0.05. *(B)* Hierarchical clustering of the time course plasma samples resolved by the DIA-SWATH-MS proteomic analysis. *(C)* Non-negative matrix factorization (NMF) analysis of the plasma proteome changes converges into three major protein signatures with distinct temporal behavior. The top graph depicts the model fit for the NMF computed at k=2-10 across 200 random initializations. The bottom heatmap represents the distribution of the three major protein signatures selected from the top scoring model (k=3). *(D)* Representative normalized intensity values for the top proteins in the NMF Signature 1, *(E)* Signature 2, and *(F)*, Signature 3. *(G)*, Schematic depiction of the two-staged model of the plasma proteome trajectory over the course of infection.

### Vascular proteome responses are temporally coordinated in an organ-specific fashion

Vascular surface proteins are located at the blood-tissue interface of the organs, which makes them potential targets of early molecular perturbations induced by blood infections. To examine whether sepsis triggers a similar staging pattern in the vascular proteome as observed in the plasma proteome, infected mice underwent systemic perfusion using Sulfo-NHS-biotin to tag proteins located on the vascular surfaces(17). This technique greatly enriches for proteins accessible to the blood, with very little penetration into the parenchyma based on streptavidin staining of tissues (17). Biotinylated proteins were enriched on streptavidin columns and analyzed by DIA-SWATH-MS. Thousands of proteins were accurately quantified across the vasculature of the organs (Fig. 2A), resulting in a substantial increase in the number of identifications compared with our previous studies using regular shotgun proteomics (SI appendix, Fig. S2, Supplementary table 1). Additionally, statistical analysis indicated that a large fraction of the vascular proteome was significantly perturbed during infection (Fig. 2B). Clustering of the data shows that the septic vascular proteome has a more complex kinetics than the plasma proteome, displaying clusters with similar temporal behaviors across the organs, and protein clusters exhibiting organ-specific changes (Fig. 2C).

**Fig. 2.**
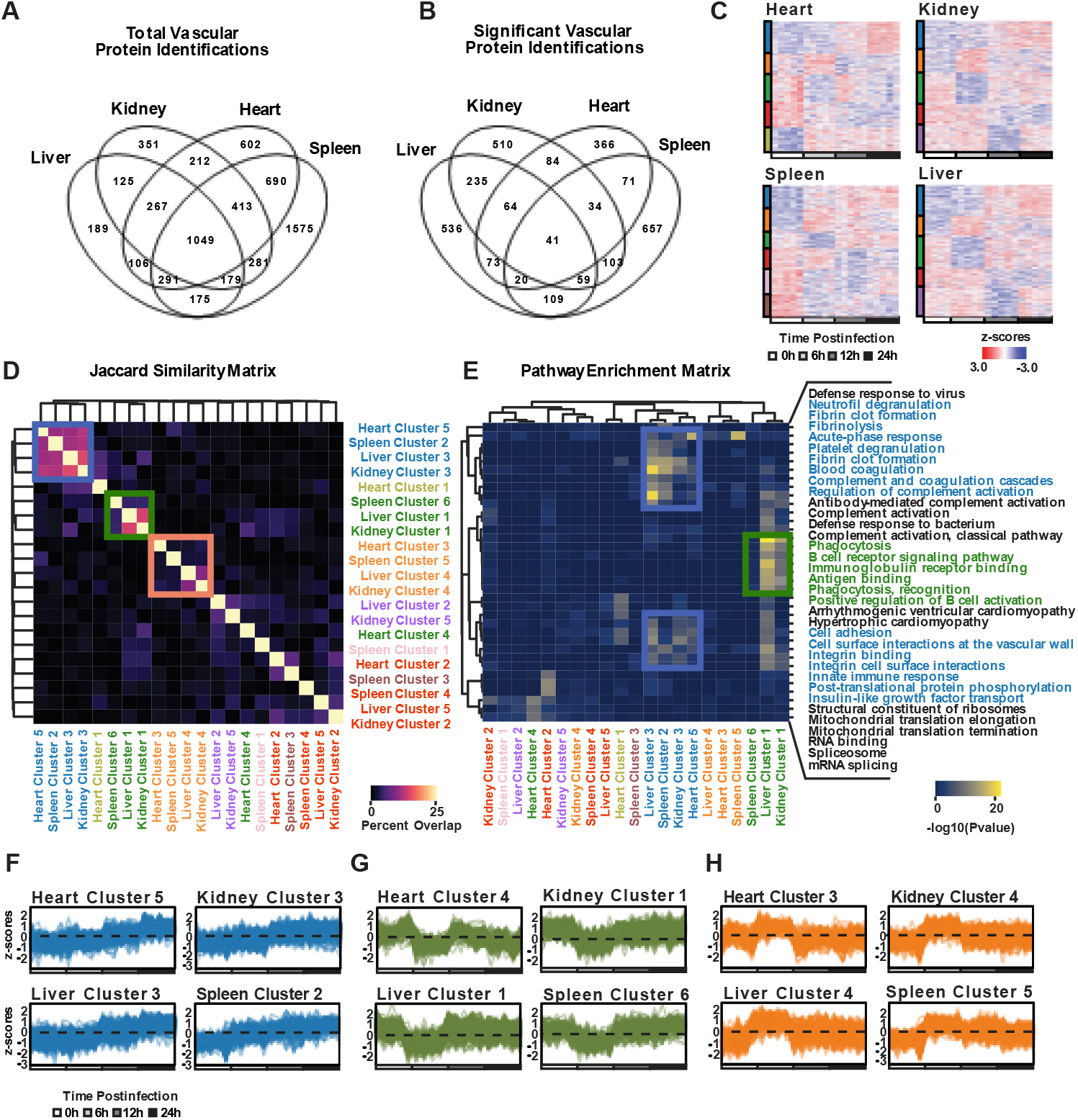
Coordinated vascular proteome responses triggered by S. aureus sepsis. *(A)* Total number of proteins identified through systemic vascular tagging across liver, kidney, heart, and spleen at 0h, 6h, 12h, and 24h postinfection (n=4-5 per group). *(B)* Significantly altered proteins using a 1-way ANOVA (FDR < 0.10) across the four organs and the intersections between them. *(C)* Heatmaps showing the k-means clustering of significant protein identifications in the heart, kidney, spleen, and liver. Clusters with the same proteome trajectory (i.e., temporal intensity patterns) are coded with the same color as represented in the bars to the left of the heatmaps. *(D)* Jaccard similarity matrix (intersection over the union) representing the percent overlap of the protein identifications across vascular clusters. Blue and green squares refer to subsets of vascular clusters displaying high overlap of proteins and the orange square, a subset of vascular clusters displaying low overlap. All cluster colors are according to the proteome trajectories. *(E)* Hierarchical clustering of the main biological pathways obtained from functional enrichment analysis of the vascular clusters colored according to P-value. *(F)* Trace representations of the normalized protein abundances (z-scores) for the vascular clusters that increase over time, *(G)* decrease at 6h or *(H)* increase at 6h post infection.

To determine if vascular proteome trajectories shared across the organs reflect the regulation of similar proteins, all vascular clusters were extracted, and both their compositions, sharedness degree and functional enrichment were evaluated. Four clusters (heart cluster-5, spleen cluster-2, liver cluster-3 and kidney cluster-3; Blue) shared a significant number of proteins (∼15%), and common biological pathways, including neutrophil and platelet degranulation, activation of coagulation and complement, and regulation of leukocyte-endothelial interactions (Fig. 2D-E). The abundance of all proteins in these clusters was linearly increased over the time of infection (Fig. 2F). Additionally, three other clusters (spleen cluster-6, liver cluster-1 and kidney cluster-1; Green) also shared many proteins (∼14%), as well as pathways related to phagocytosis and the adaptive immune response, particularly receptor mediated interactions (Fig. 2E and 2G). Interestingly, the abundance of all proteins in these clusters dramatically dropped at 6h post infection, suggesting the rapid downregulation and/or shedding of a large fraction of the vascular proteome. In sharp contrast to these general responses across the organs, evidence of tissue-specific changes was also detected. For example, some specific clusters (heart cluster-3, spleen cluster-5, liver cluster-4 and kidney cluster-4; Orange) shared only a few proteins (∼2%), suggesting the induction of organotypic responses (Fig. 2D). Functional enrichment linked these clusters to ion transport and homeostasis (heart cluster-3), metal sequestration by antimicrobial proteins (spleen cluster-5), cellular responses to labile heme (liver cluster-4) and mitochondrial dysfunction (kidney cluster-4). Notably, despite the marked differences in their protein contents, these clusters exhibited synchronous proteome trajectories (Fig. 2H). Taken together, vascular responses to sepsis entail both general and organotypic alterations of the vascular proteome very early during disease progression.

### Comparison between parenchymal and vascular proteome alterations during sepsis

To determine the progression of parenchymal proteome responses, infected organs were subjected to whole tissue proteomics analysis, which resulted in the accurate quantification of thousands of proteins (Supplementary table 2). Like the vascular proteome, parenchymal responses displayed both shared and organ-specific proteome trajectories (SI appendix, Fig. S3). Principal component analysis (PCA) shows that parenchymal proteome responses were highly heterogeneous across the organs, displaying different temporal patterns (Fig. 3A-D). For example, the liver tissue was governed by early changes and a significant separation of the samples at 0h and 6h (Fig. 3A), whereas the heart exhibited a more delayed response (no notable difference between 0h and 6h, but clear differences between samples at 6h and 12h) (Fig. 3B). The kidney and the spleen were also quick responders, although the spleen showed the largest parenchymal proteome changes of all organs, with >2000 proteins significantly altered during sepsis (Fig. 3C-D). Interestingly, the analysis of the vascular proteomes shows a rather uniform pattern (Fig. 3E-H), in contrast to the heterogeneity of the temporal patterns of the parenchymal responses. All vascular proteome datasets displayed a distinct separation of the 0h and 6h time points, and clustering of the 12h and 24h time points, independently of the organ. Despite this similar behavior, the number of dysregulated proteins that were shared across the organ vascular fractions at 6h was very low (41, ∼1.4%). These findings suggest that sepsis induces early changes in the vascular proteome of all organs, but each proteome response is most likely shaped by organ specific factors.

**Fig. 3.**
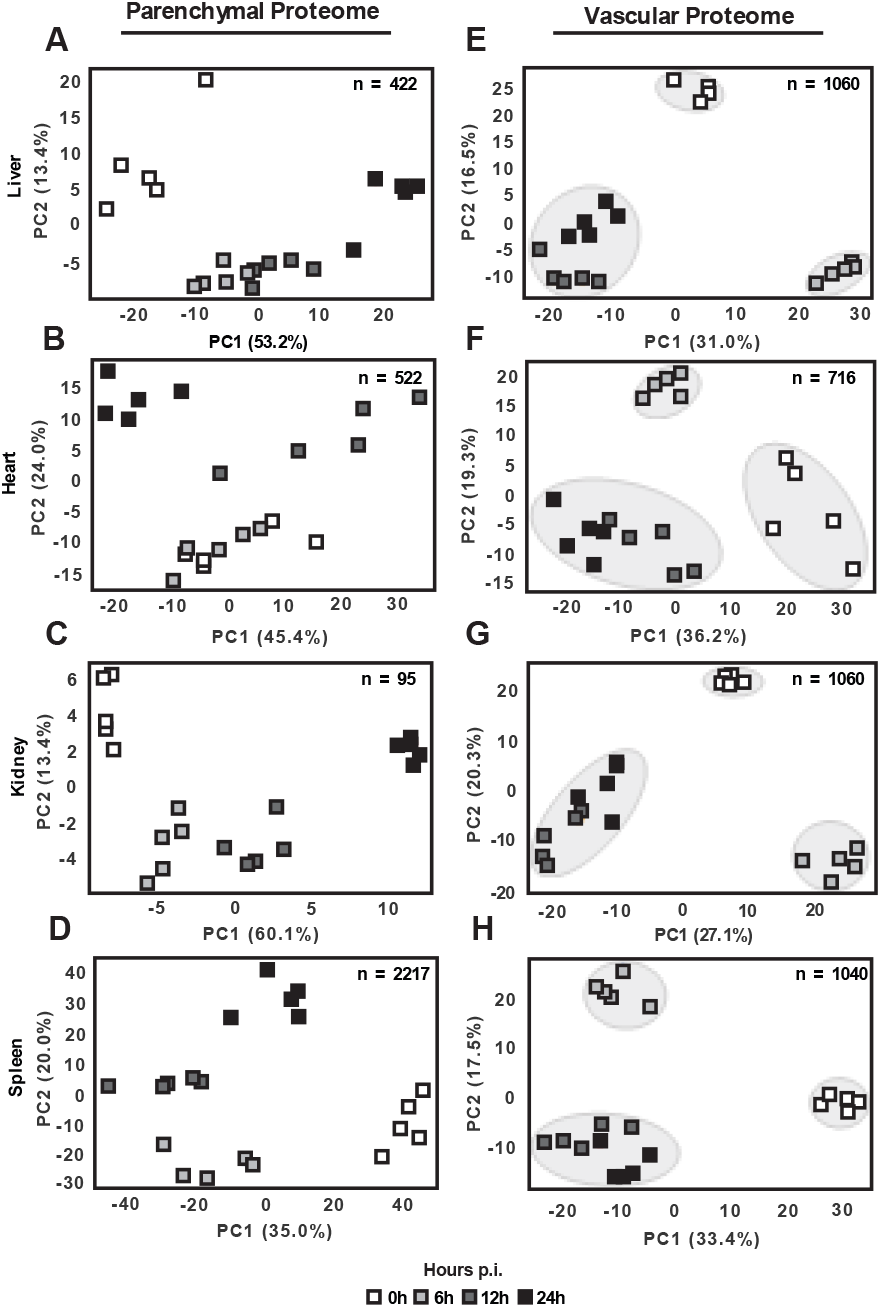
Principal component analysis (PCA) of significantly altered parenchymal and vascular proteins during infection. *(A)* Proteins identified displaying statistically significant temporal patterns using a 1-way ANOVA (FDR < 0.10) were subject to PCA analysis to identify time dependent variability for liver parenchymal proteins (n=422), *(B)* heart parenchymal proteins (n=522), *(C)* kidney parenchymal proteins (n=95), *(D)* spleen parenchymal proteins (n=2217), *(E)* liver vascular proteins (n=1060), *(F)* heart vascular proteins (n=716), *(G)* kidney vascular proteins (n=1060), and *(H)* spleen vascular proteins (n=1040).

### Building the proteome trajectory of sepsis through a higher-order networking approach

To reconstruct the disease trajectory of sepsis using the host proteome response as a readout, we developed a novel computational pipeline to integrate the data across time points, organs, and tissue compartments. Correlative Stratification of Proteome Trajectories (COR-SPOTS) was designed as a higher-order networking approach to store, organize and integrate multiple data layers based on temporal and functional relationships between the data points (see material and methods). Significantly dysregulated proteins from each organ and compartment (plasma, vascular, and parenchymal) were parsed through STRING-DB to group proteins based on functional associations(22). Highly interconnected clusters of nodes were then identified through the Louvain algorithm for community detection, and collapsed into new single nodes. We then computed ∼24 million pairwise spearman correlations over all combinations of detected community nodes, which resulted in ∼3.7 million significant correlations (false discovery rate; FDR<0.10) that were used to define the linking edges of the network. Approximately an equal number of significant positive and negative correlations were observed. Other similarity relationships between the nodes such as the content overlap and the functional enrichment were also stored within the network, resulting in a compiled and fully searchable small database containing all proteomics results and metadata from this study, which can be found on the online repository the Network Data Exchange (NDEx) with UUID: 45474980-9d56-11eb-9e72-0ac135e8bacf (Fig. 4A).

**Fig. 4.**
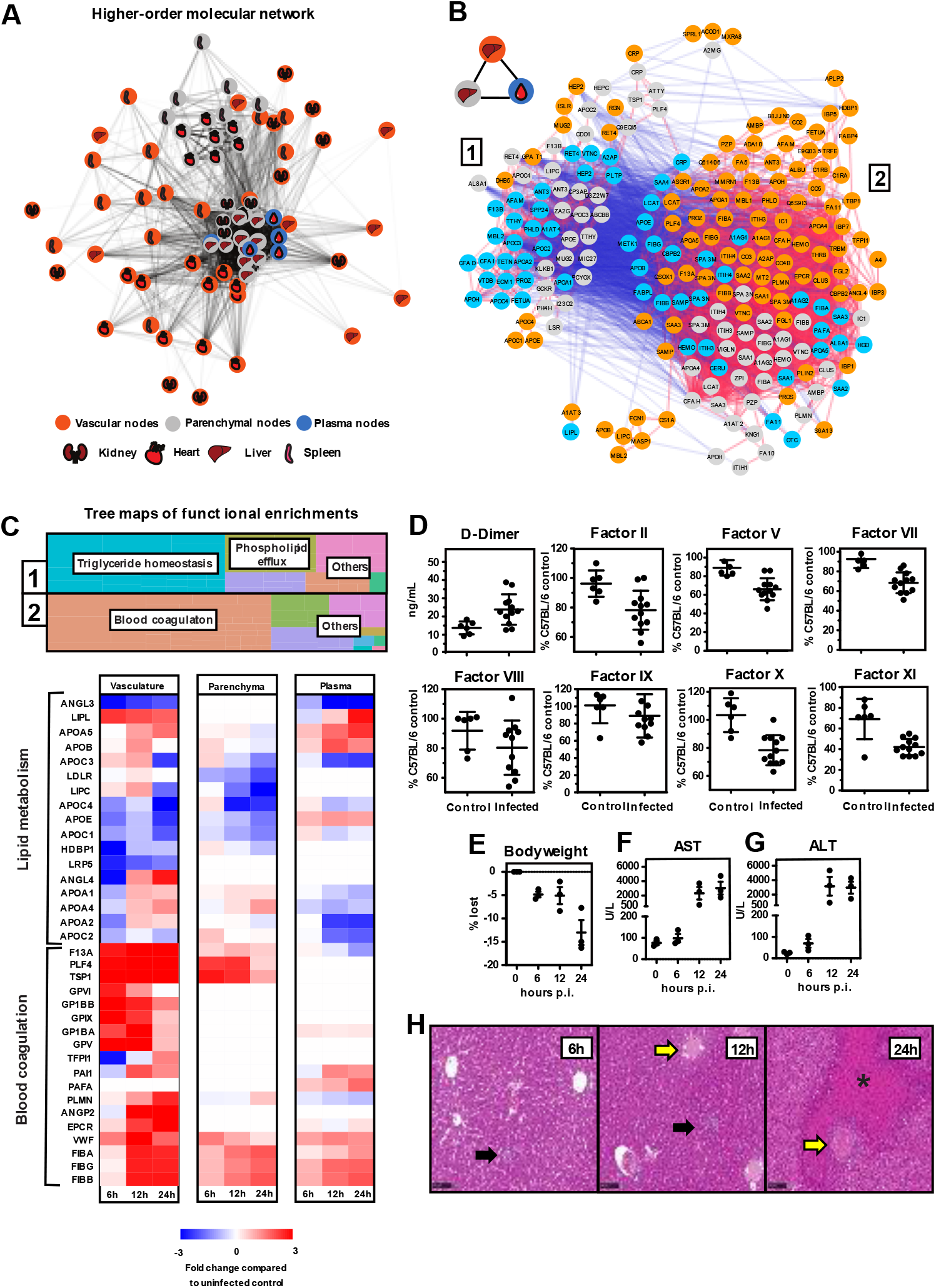
Higher-order molecular networking reveals vascular proteome perturbations preceding coagulopathy and hepatic damage. *(A)* Complete network analysis of all proteome changes across organs and tissue compartments visualized by the edge-weighted spring embedded layout, where the edge distance and shade represent the proportion of significant pairwise correlations between two nodes. Each node color represents the compartment of origin with tissue represented by grey fill color, vascular represented by orange fill color, and plasma represented by blue fill color. *(B)* Functional crosstalk between liver parenchymal, vascular, and plasma proteins. Three intercorrelated higher-order nodes were selected and their pairwise protein correlations were extracted. The subnetwork contains 2 major anti-correlated clusters of proteins. Edges are colored using a blue, white, red continuous scale ranging from -1.0 to 1.0. *(C)* Functional enrichment analysis of the 2 major protein clusters visualized by treemaps (Top). Perturbation of lipid metabolic proteins and upregulation of blood coagulation components represented by compartmentalized heatmaps of the liver parenchymal tissue, vasculature, and blood plasma. Log2 fold change values are compared to uninfected proteome measurements (Bottom). *(D)* Enzyme activity assays measuring blood coagulation factors present in murine plasma at 24h post infection. *(E)* Changes in body weight, *(F)* blood glucose and the markers of hepatic damage *(G)* AST and *(H)* ALT. *(I)* Representative images of the hematoxylin and eosin stain of liver tissue over the time course. Leukocyte infiltration was noticed as early as 6h post infection (black arrows) and vascular occlusion and thrombosis as early as 12h post infection (yellow arrows), necrotic areas are marked with an asterisk (*).

### Vascular proteome responses in the liver precede coagulopathy and hepatic damage

We have previously shown that the murine model of *S. aureus* sepsis suffers from a dominant coagulopathic phenotype in the liver(17). To demonstrate the utility of COR-SPOTS to dissect molecular changes across the blood-tissue interfaces, we ranked the nodes and selected the hepatic vascular, parenchymal and plasma nodes with the highest inter-correlation. Proteins stored within these highly correlated nodes displayed a bimodal network topology characterized by the presence of two distinct anti-correlated clusters of proteins (Fig. 4B). Cluster 1 contained most of the plasma and parenchymal proteins linked to triglyceride and phospholipid metabolism, whereas cluster 2 was enriched in vascular proteins linked to coagulation (Fig. 4C). Many proteins involved in lipid metabolism were rapidly diminished from the hepatic vasculature, including multiple apolipoproteins, angiopoietin-like proteins, and the glycosylphosphatidylinositol-anchored high-density lipoprotein-binding protein 1 (HDBP1). Some of these factors were also reduced in the liver parenchyma and plasma during infection, but these changes were more delayed compared with the vascular proteome. On the other hand, proteins involved in coagulation were rapidly accumulated in the hepatic vasculature. This included deposition of several platelet glycoproteins (GPIX, GP1BB, GP1BA, GPV), indicating abundant intravascular platelet deposition, together with an increase in the levels of Von Willebrand Factor (VWF) and fibrinogen. Most of these changes were specific to the vascular proteome and were only weakly captured in the parenchyma and plasma compartments towards the later time points (12h and 24h). Notably, the levels of Tissue Factor Pathway Inhibitor 1 (TFPI1) at 6h post infection were rapidly depleted, strongly suggesting a prothrombotic priming of the liver vasculature very early during infection (Fig. 4C). Further confirming that *S. aureus* bacteremia results in systemic coagulopathy, the levels of D-dimers in plasma were significantly increased at 24h post infection, and widespread consumption of multiple coagulation factors was also confirmed by enzyme activity assays (Fig. 4D). Metabolic changes were also detected since the mice lost significant amounts of weight over time (Fig. 4E). Interestingly, organ failure based on the increased levels of ALT and AST in circulation was not apparent until 12h post infection (Fig. 4F-G), which coincided with the occurrence of massive thrombosis and tissue necrosis in the liver towards the late time points (Fig. 4H). Taken together, the data suggest a global reprogramming of the liver by shutting down metabolic functions as the level of intravascular coagulation and vasculopathy increases. More importantly, these global changes were preceded by early alterations of the vascular proteome, long before signs of organ failure were detected through well-established correlates of organ damage.

### Vascular proteome responses in the heart precede bacterial invasion and neutrophil infiltration

Comparison between the changes induced by sepsis in the vascular and whole organ proteomes indicated a delayed response of the cardiac parenchyma compared with the rest of the organs (Fig. 3B). Inspection of the cardiac nodes in the network showed a distinct separation of the parenchymal nodes from the vascular nodes, distinguishing the heart from the other organ proteomes (Fig. 4A). We hypothesize that this disparity in the cardiac protein response was linked to a difference in the kinetics of the disease trajectory of sepsis in the heart, and perhaps linked to the presence or absence of major cellular events, such as bacterial invasion and/or leukocyte infiltration of the organs. The bacterial burden of the organs was in line with this hypothesis, showing that despite the presence of bacteria in the liver, kidney, and spleen already by 6h post infection, no bacteria were found in the heart until 12h after inoculation (Fig. 5A). Since neutrophils are the primary responders to the presence of bacterial infiltrates in the tissues, we assessed the levels of myeloperoxidase (MPO) and neutrophil elastase (ELANE), two granule proteins that are specific to neutrophil degranulation and formation of extracellular traps. The levels of MPO and ELANE in the liver were rapidly raised upon infection both in the parenchyma and vascular proteomes, whereas their deposition in the cardiac parenchyma and vasculature was first observed after 12h (Fig. 5B). Histological analysis confirmed that neither bacterial microabscesses nor immune infiltrates were observable in heart tissue until 12h post infection (Fig. 5C). Taken together, these data suggests that in this model of *S. aureus* sepsis, bacterial dissemination and leukocyte infiltration are temporally delayed in the heart compared to the other organs, opening an interesting window to capture very early molecular changes in the vasculature as a response to systemic inflammation.

**Fig. 5.**
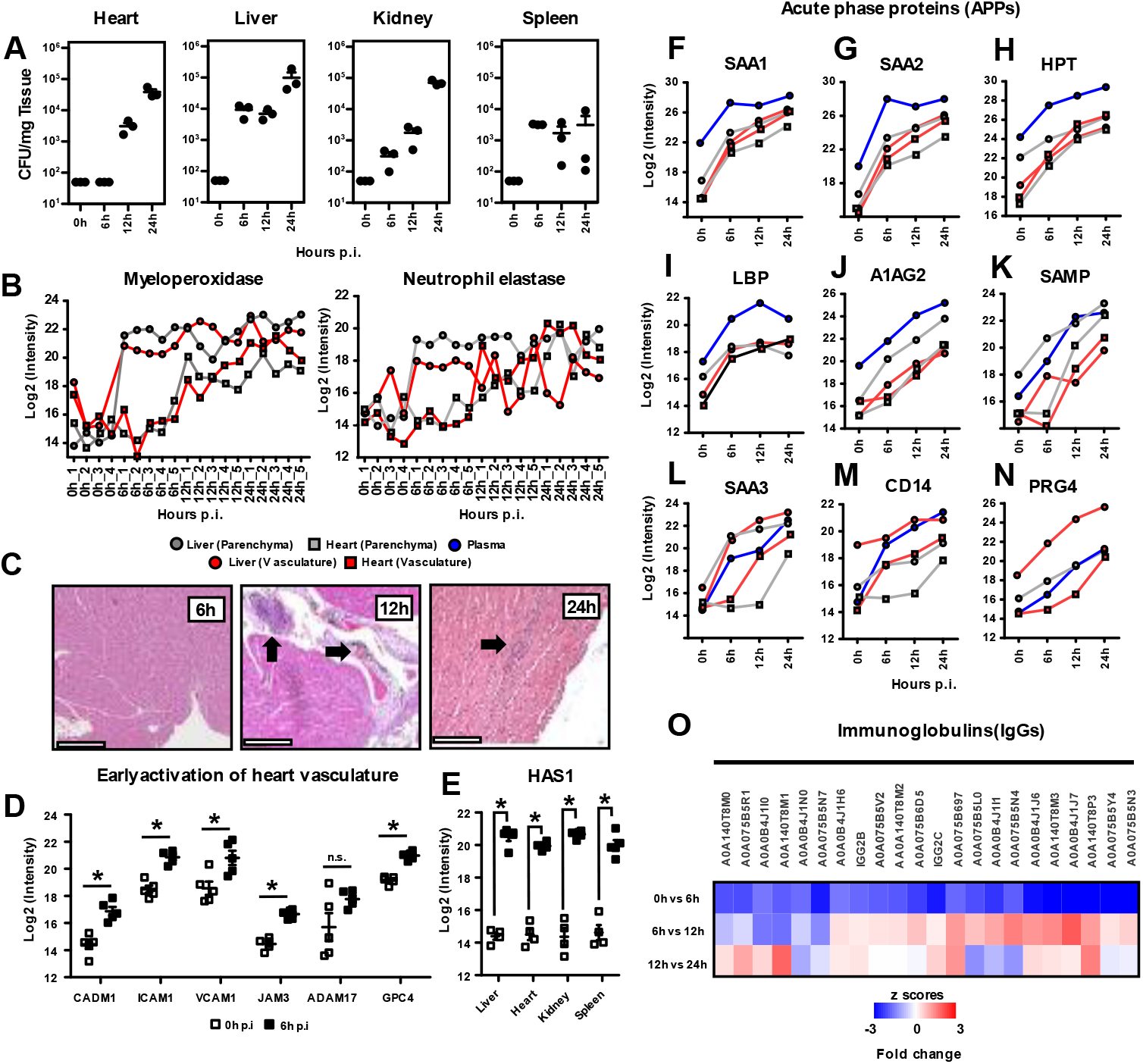
Bacterial invasion and neutrophil infiltration are delayed in the heart when compared to other organs. *(A)* Organs were harvested, homogenized, and plated to quantify the bacterial colony forming units (CFU) present over the experiment time course. *(B)* Proteomic measurements of myeloperoxidase (MPO) and neutrophil elastase (ELANE) for both liver and heart vascular and parenchymal proteomes. Liver measurements are represented by filled circles; heart measurements are represented by filled squares. Tissue was represented by grey fill color, and vascular measurements by orange fill color. Plasma measurements were represented by filled blue circles. *(C)* Representative images of histological analysis of heart tissue over the experiment time course. Bacterial and leukocyte infiltration was first noticed at 12h and 24h post infection (black arrows). *(D)* Cardiac vascular proteins changing very early during infection. *(E)* Hyaluronan Synthase 1 (HAS1) was significantly increased in the vascular surface of all four organs. *(F-N)* Global vs compartmentalized changes in the levels of acute phase reactants (APRs) in liver, heart and plasma. Plot labeling as described previously. *(O)* 23 immunoglobulin proteins found to dramatically decrease in abundance as early as 6h post infection. Log2 fold change values are compared to the previous time point.

Next, we focused on the changes in the heart vasculature at 6h to identify molecular alterations that precede bacterial and leukocyte invasion of the tissue (Fig. 5D). Multiple vascular adhesion proteins were rapidly induced during infection, such as CADM1, ICAM1 and VCAM1. Upregulation of junctional endothelial proteins such as JAM3 was also observed, together with an increase in the levels of ADAM17, a protease that has been linked to most of the early shedding of the vascular surfaces upon inflammation. Changes in the glycocalyx were also noticed, including upregulation of heparan sulfate proteoglycans such as GPC4, and very robust induction of the hyaluronic acid synthase HAS1. In fact, early upregulation of HAS1 was consistently observed across all vascular surfaces, directly implicating HAS1 in glycocalyx remodeling during sepsis (Fig. 5E).

### Compartmentalization of acute-phase reactants and immunoglobulin shedding are early vascular responses to *S. aureus* sepsis

Acute phase reactants (APRs) are inflammation markers that exhibit rapid and substantial changes in serum concentration during inflammation. Most APRs are synthesized in the liver as a response to both local and systemic inflammation, and some of them get associated with the endothelium, where they exert multiple functions. We used COR-SPOTS to map the dynamics of APRs during infection across organs and compartments. Although the dynamics of the APRs in plasma followed a linear increase, as has been previously described, when mapping the levels of APRs associated with parenchymal and vascular compartments, we noticed at least two distinct patterns. Type I APRs experienced rapid induction in all organs and compartments, and steadily accumulated over time. This pattern was characteristic of APRs such as SAA1, SAA2, HPT and LBP (Fig. 5F-I). However, we also identified a second group or Type II APRs, whose levels were compartmentalized and sensitive to the disease status of the organs. For example, A1AG2 and SAMP levels in liver parenchyma and vascular fractions followed the same kinetics as in plasma (Fig. 5J-K). Interestingly, their levels in cardiac parenchyma and vasculature remained unchanged at 6h, followed by a linear increase at 12h and 24h. Other APRs such as SAA3 followed a similar trend, except for a temporal difference between the cardiac vascular proteome response (up at 12h) and the cardiac parenchymal response (up at 24h). CD14 was also increased in the liver and in the heart vasculature by 6h, but increased cardiac parenchymal levels were only detected by 24h. We have previously shown that PRG4, a protein that behaves as an APR in many inflammatory models gets deposited in the liver vasculature in this sepsis model of *S. aureus* bacteremia by 24h. Looking at the dynamics of PRG4 across organs and compartments, it became apparent that vascular accumulation starts early in the liver and increases linearly, but association with the heart vasculature could not be detected until the late time points. Finally, we also observed that in contrast to the increasing levels of many vascular adhesion molecules and APRs, the most dramatic downregulation taken place in the cardiac vasculature at 6h was the shedding of many immunoglobulins (IgGs), probably loosely associated with the vascular wall, without changing IgG total plasma levels. All these changes are consistent with a massive remodeling of the vascular proteome during sepsis, even before bacteria or infiltrating immune cells are observed. These changes are expected to significantly alter the physicochemical properties of the vascular surfaces, and their organotypic responses to systemic infection.

## Discussion

Understanding the relationships between vascular failure and organ damage has important implications for sepsis therapeutics and diagnostics. Even though many patients quickly respond to standard care, an unacceptably high fraction of cases still progress into intractable shock responses with a high risk for long term morbidity and deadly outcomes. Vascular stabilization as a therapeutic goal in sepsis has been discussed for years, although few preclinical studies have been conducted to assess the impact of vasculoprotective approaches on relevant sepsis outcomes, and so far, none of them have succeeded in making a translational jump from the bench to the bedside. Part of the problem lies on the enormous heterogeneity of the organ vasculatures, and a lack of detailed knowledge on the molecular composition of the vascular surfaces and how they are affected by sepsis.

Here, we demonstrate that sepsis triggers a series of well-coordinated proteome changes across the blood-tissue interfaces of the organs, but the nature of those changes and their specific temporal patterns are largely organotypic. These organ-specific patterns seem to be partially linked to the occurrence of major sepsis “checkpoints”, such as systemic vascular activation or the actual invasion of the organs by pathogens and infiltrating immune cells. Interestingly, the liver undergoes a rapid reprogramming of its vascular proteome by downregulating lipid metabolic functions and becoming primed towards a prothrombotic and anti-fibrinolytic state, long before signs of organ dysfunction and actual thrombosis are observable. This finding is remarkable, firstly because it suggests that vascular proteome alterations can have a yet unrealized potential as a source of new generation diagnostic and prognostic markers. Secondly, it also opens the possibility that timely therapeutic interventions targeting the vasculature might prevent irreparable organ damage. For example, we observed that hepatic thrombosis is preceded by early depletion of vascular TFPI, upregulation of VWF and intravascular platelet deposition. Alterations in the extrinsic pathway of coagulation were also reported in baboon models of gram-negative sepsis, and exogenous TFPI administration rescues the organs from coagulation-induced damage(23, 24). However, there can be difficulties in translating some of the findings from preclinical models to therapy(25, 26). For example, results from clinical trials to evaluate the efficacy and safety of Tifacogin, (recombinant TFPI) to treat sepsis have so far been disappointing, emphasizing the complexity of the host response to systemic infection.

The host response to sepsis is overwhelmingly complex since it encompasses crosstalk between multiple systems, and a time-dependent staging, where both pro- and anti-inflammatory components might be operating at different time points. Predicting the disease trajectory of sepsis is therefore a most sought-after clinical goal. Here we show that the host proteome response to sepsis is a powerful molecular readout of the septic response. Using a novel networking approach, COR-SPOTS, we were able to identify very early vascular changes that even precede bacterial and leukocyte infiltration of the tissue. Some of these responses have been previously reported, such as the upregulation of vascular adhesion molecules and glycocalyx components, whereas other were completely unexpected, such as the compartmentalization of the acute-phase reaction and the massive shedding of vasculature-associated IgGs. Future work should explore how these early vascular events are coordinated, for example by focusing on transcriptional and/or posttranslational mechanisms of proteome regulation.

Finally, the accumulating data suggests that sepsis entails a series of heterogeneous responses to systemic infection, with distinct molecular trajectories and disparate clinical outcomes. The networking strategy deployed by COR-SPOTS can be exploited to construct concise molecular readouts of the host response, and to compare the impact of individual pathogens and/or virulence factors to the molecular heterogeneity of sepsis. Similarly, our approach might also facilitate the quick evaluation of theragnostic markers that are sensitive to treatment options, in order to find novel indicators that reflect the status of critically ill patients and to accelerate the process of clinical decision-making in intensive care units.

## Materials and Methods

### Bacterial preparation and infection

*S. aureus* strain USA300/TCH1516 was grown at 37 °C in liquid cultures of Todd-Hewitt broth (THB, Difco) with agitation (200 rpm), and later incubated in 5 mL of fresh THB overnight. Roughly, 400μL of the overnight culture was inoculated into 6 mL of fresh THB and incubated until an OD600 = 0.4 was reached. Bacteria were centrifuged, washed twice, and resuspended in PBS. 8–10-week-old C57Bl/6 mice were intravenously infected through the retroorbital sinus with 5 × 10^7^ CFU of the bacteria culture in 100μL PBS, or just with 100μL PBS in the control group. Animals were euthanized at 6h, 12h or 24h using isoflurane. One group of animals was immediately subjected to chemical biotinylation perfusions as described below, and another group was subjected to cardiac puncture to collect blood samples. After these procedures, multiple organs were collected from both groups (liver, kidney, heart and spleen). All animals were housed in individual ventilated cages in vivaria approved by the Association for Assessment and Accreditation of Laboratory Animal Care at the School of Medicine, UC San Diego. All experiments followed relevant guidelines and regulations consistent with standards and procedures approved by the UC San Diego Institutional Animal Care and Use Committee (protocol #S99127 and #S00227M).

### Systemic chemical perfusions

Chemical perfusions were performed as previously reported(17). Briefly, mice were subjected to a median sternotomy and the left ventricle of the heart was punctured with a 25-gauge butterfly needle (BD Vacutainer). A small cut was made in the right atrium to allow draining of perfusion solutions. All ice-cold perfusion reagents were infused using a perfusion pump (Fischer scientific). Blood was washed out with PBS for 5 min at a rate of 5 mL/min. A solution containing 100mM EZ-link Sulfo-NHS-biotin (Thermo Fischer) in PBS, pH 7.4 was perfused at a rate of 3 ml/min for 10 min. Finally, a quenching solution (50mM Tris-HCl, pH 7.4) was perfused at 3 ml/min for 5 min. Control animals were perfused with PBS only.

### Organ preparations

Collected organs were homogenized in a buffer containing 5M urea, 0.25M NaCl and 0.1% SDS. Samples were briefly centrifuged for 5 min and the supernatant was transferred to new tubes. Protein was quantified by standard BCA assay (Thermo Scientific) as per manufacturer instructions. Samples were stored at −80 °C until further analysis. In parallel, a piece of each organ was homogenized in 1 mL ice-cold PBS, and the samples were plated on Todd-Hewitt Agar for CFU analysis.

### Blood Chemistry and Coagulation Assays

Whole blood was collected via cardiac puncture and placed in a pro-coagulant serum tube (BD Microtainer #365967) for 4 hours at room temperature. Serum was isolated by spinning the tubes at 2000xg and collecting the supernatant. All samples were frozen and thawed once before analysis. Blood chemistry parameters were measured on a Cobas 8000 automated chemistry analyzer (Roche) with a general coefficient of variance of <5%. All samples were frozen and thawed no more than two times before analysis. Blood coagulation factor assays were performed as previously described (27).

### Bacterial Colony Forming Units Counts

Organs of interest were harvested at the indicated time points and placed in a 2mL tube (Sarstedt #72.693.005) containing 1 mL ice cold PBS and 1.0 mm diameter Zirconia/Silica beads (Biospec Products #11079110z). Samples were homogenized using a MagNA Lyzer (Roche) for 2 minutes at 6000 rpm. Whole blood was collected via cardiac puncture and placed in an EDTA tube (BD Microtainer #365974). An aliquot of each organ or blood sample was serially diluted in PBS and plated on Todd-Hewitt Agar to enumerate CFU.

### Histological Analysis

Tissues were harvested at the indicated time points and fixed in 10% buffered formalin (Fischer Chemical) for 24 hr, followed by submersion in 70% ethanol for at least 24hr. The samples were paraffin embedded, sectioned (3 μm), and stained with hematoxylin/eosin.

### Purification of biotinylated proteins

Biotinylated proteins were isolated from the organ homogenates using a Bravo AssayMap platform and AssayMap streptavidin cartridges (Agilent). The cartridges were equilibrated with ammonium bicarbonate (50mM, pH 8), and biotinylated samples were loaded. Non-biotinylated proteins were removed by extensive wash with 8M urea in 50 mM ammonium bicarbonate buffer (pH 8). Cartridges were further washed with Rapid digestion buffer (Promega, Rapid digestion buffer kit) and bound proteins were subjected to on-column digestion using mass spec grade Trypsin/Lys-C Rapid digestion enzyme (Promega, Madison, WI) at 70 °C for 2 h. Released peptides were desalted using AssayMap C18 cartridges (Agilent). Samples were stored at −20 °C prior to DIA-SWATH-MS analysis

### DIA-SWATH-MS

DIA-SWATH-MS analysis was performed on a Q Exactive HF-X mass spectrometer (Thermo Fisher Scientific) coupled to an EASY-nLC 1200 ultra-high-performance liquid chromatography system (Thermo Fisher Scientific). Peptides were trapped on a pre-column (PepMap100 C18 3 μm; 75μm x 2cm, Thermo Fisher Scientific) and separated on an EASY-Spray column (ES803, column temperature 45 °C, Thermo Fisher Scientific). Equilibrations of columns and sample loading were performed as per manufacturer’s guidelines. Solvent A (0.1 % formic acid), and solvent B (0.1 % formic acid, 80% acetonitrile) were used to run a linear gradient from 5 % to 38 % over 120 min at a flow rate of 350 nl/min. A DIA method was implemented using a schedule of 44 variable acquisition windows as previously reported (19). The mass range for MS1 was 350-1,650 m/z with a resolution of 120,000 and a resolution of 30,000 for MS2 with a stepped NCE of 25.5, 27 and 30.

### DIA-SWATH-MS data analysis

An in silico spectral library was generated for the reference Mus musculus proteome (EMBL-EBI RELEASE 2020_04: 22,295 entries) using deep neural networks as implemented in DIA-NN (v1.7.10)(20). For search space reduction a list of previously MS detectible mouse peptides were compiled from the Peptide Atlas Project (28), and from two published large-scale DIA proteomics studies (11, 29). The compiled library with a final number of 667,455 precursors was used for DIA data extraction with a protein q-value of 0.01 and RT-profiling enabled.

### Statistical Analysis

All statistical methods were implemented using Python 3.6.10. Proteomics results from the DIA-SWATH-MS analysis were filtered using a one-way ANOVA followed by a Benjamini–Hochberg procedure to control for a false discovery rate (FDR) < 0.10. Statistically significant identifications were further subjected to principal component analysis (PCA). Proteins were given a standardized score using a z-score normalization. Proteomics results were separately analyzed using Welch’s t-test to generate volcano plots and heatmaps.

### Functional Enrichment Analysis

Functional enrichment analysis of differentially abundant proteins was performed through the Database for Annotation, Visualization, and Integrated Discovery (DAVID). DAVID was run using default settings with the thresholds count ≥ 2 and EASE ≤ 0.1 (30). Visualization of enriched Genome Ontology (GO) terms was produced using code adapted for treemap visualizations from the Web tool REVIGO(31).

### Clustering

#### Non-negative matrix factorization

Plasma DIA proteomics was analyzed using non-negative matrix factorization (NMF). Following pseudocode of Lee and Seung’s multiplicative update rule, the distance between the dataset matrix (D) and the dot product of hypothetical partitions (W and H) was minimized using a coordinate descent algorithm (21). The two resulting matrices were constrained during the optimization to allow for better interpretability. First, the predictions in the matrix H must fall within the range 0.0 to the maximum observed log2(relative abundance). Second, the distributions in the W matrix must fall in the range 0.0 to 1.0 and must sum across signatures to 1.0. In order to determine the number of predicted proteome signatures, k, for each value k, 200 random initializations were made. The average Pearson correlation between the ground truth hierarchy learned from D and the weights hierarchy learned from W and average distance scores (||D -WH||F) are plotted for visual inspections at each k. Cluster number k was finally selected using the elbow selection method as well as meaningful investigation of correlation and proteome signature content.

#### Abundance Clustering

Significant proteins were grouped based on their relative abundances using k-means clustering. K selection was done through the visualization of distances to cluster centers across k values up to 20 and selecting based on the elbow method (32).

#### Functional Clustering

Protein identifications were parsed through the medium confidence (PPA>0.4) STRING-DB to build preliminary networks based on physical or functional associations. These networks were subjected to the Louvain method for community detection, a modularity based clustering algorithm. The Louvain clustering was implemented using the Python packages NetworkX and python-Louvain with default settings (33). To circumvent the heuristic property of the Louvain algorithm, ∼1000 parallel random starts were initially performed. Using the combined results from these 1000 iterations, a matrix of cluster clarity was built to store the average number of times two proteins appear in the same community, divided by the number of iterations. Using this community clarity matrix, a k-means clustering was finally performed with k being the average number of communities found across the 1000 runs.

#### Cluster Annotations

The Jaccard similarity index was also used to determine the overlap of proteins contained between two clusters by quantifying the intersection over the union. A ranked list of functional associations for each cluster were defined by using a hypergeometric test (SciPy v1.4.1). Using the Tau-b statistic from the Kendall rank correlation test (SciPy v1.4.1), functional terms between two clusters were compared by ranked enriched terms across GO terms, Reactome, and KEGG pathways.

### Higher-order Molecular Networking

All significant protein identifications from all datasets (i.e., different time points, organs, and compartments) were parsed through the functional clustering pipeline described in the previous section. Each of these higher-order community nodes contained a variable number of proteins (ranging from just a few to hundreds of proteins), operating within similar biological processes. Finally, we computed pairwise spearman correlations over all combinations of detected community nodes, which resulted in detection of significant correlations that were used to define linking edges between the nodes. Benjamini–Hochberg procedure was used to control for a false discovery rate (FDR) < 0.10. The multi-edge was then constructed using three pieces of relational information. Contained in the multi-edge are the scores: the proportion of significant pairwise spearman correlations, the Jaccard similarity index, and the Tau-b statistics previously defined. Visualization and analysis of the network layers were conducted through Cytoscape (34).

## Code Availability

We provide the code library in Python described in this work through Github: https://github.com/LewisLabUCSD/CORSPOTS. We provide jupyter notebooks in python to generate our figures and analysis.

## Author Contributions

JTS, JDE, and AGT conceived the project. GJG, CM, CP, ARC, CK and AGT conducted the mouse infection, perfusion, organ isolation, and mass spectrometry experiments. CK constructed the in-silico library for DIA-SWATH-MS and analyzed the DIA data. JTS developed COR-SPOTS. JTS, VN, ARC, JWS, JM, NEL, JDE and AGT interpreted the data. JTS and AGT prepared the figures. JTS, JDE and AGT wrote the manuscript with significant input from all authors.

## Supplemental figure legends

**Fig. S1.**
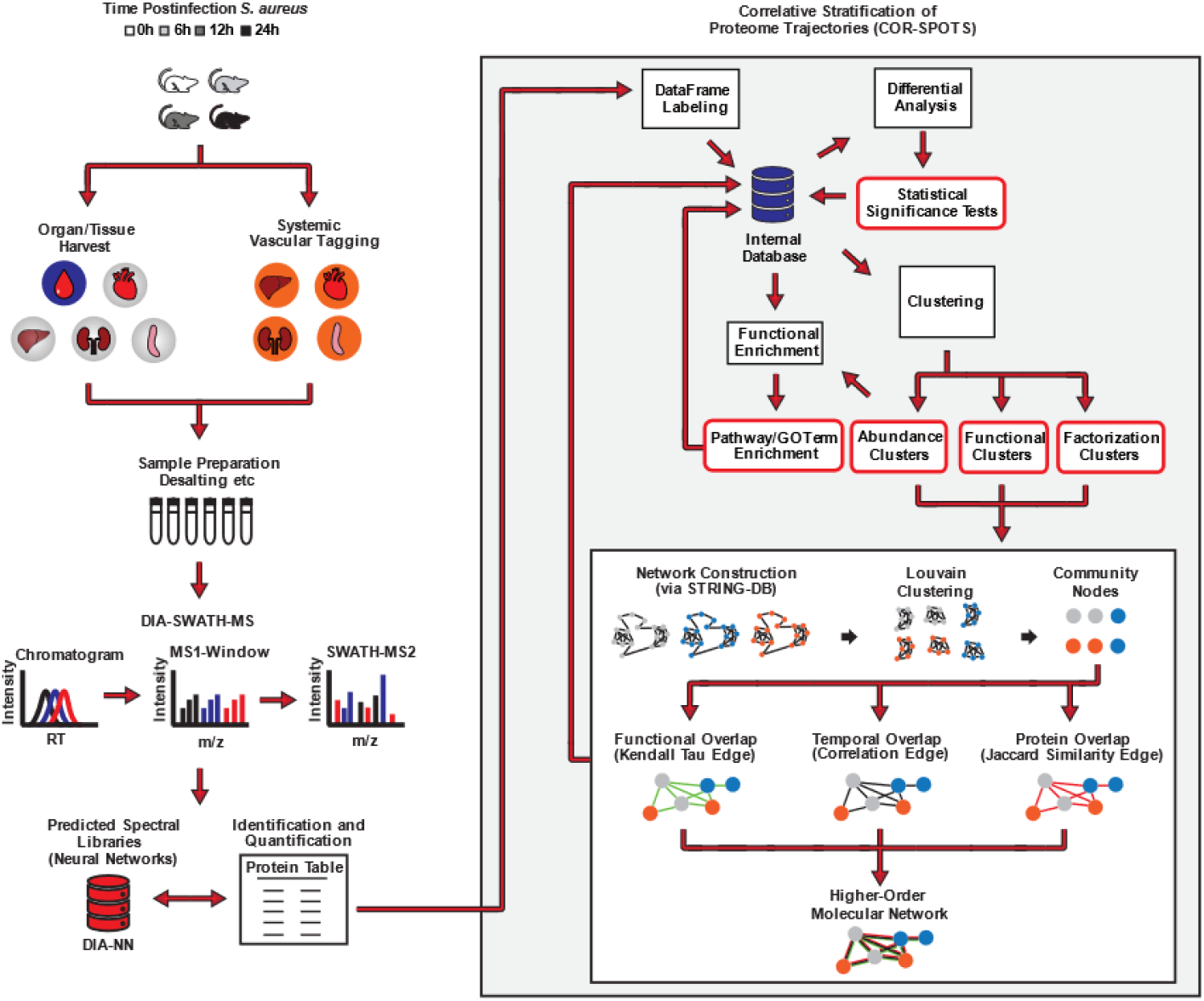
Experimental design and analytical workflow employed in the current study.

**Fig. S2.**
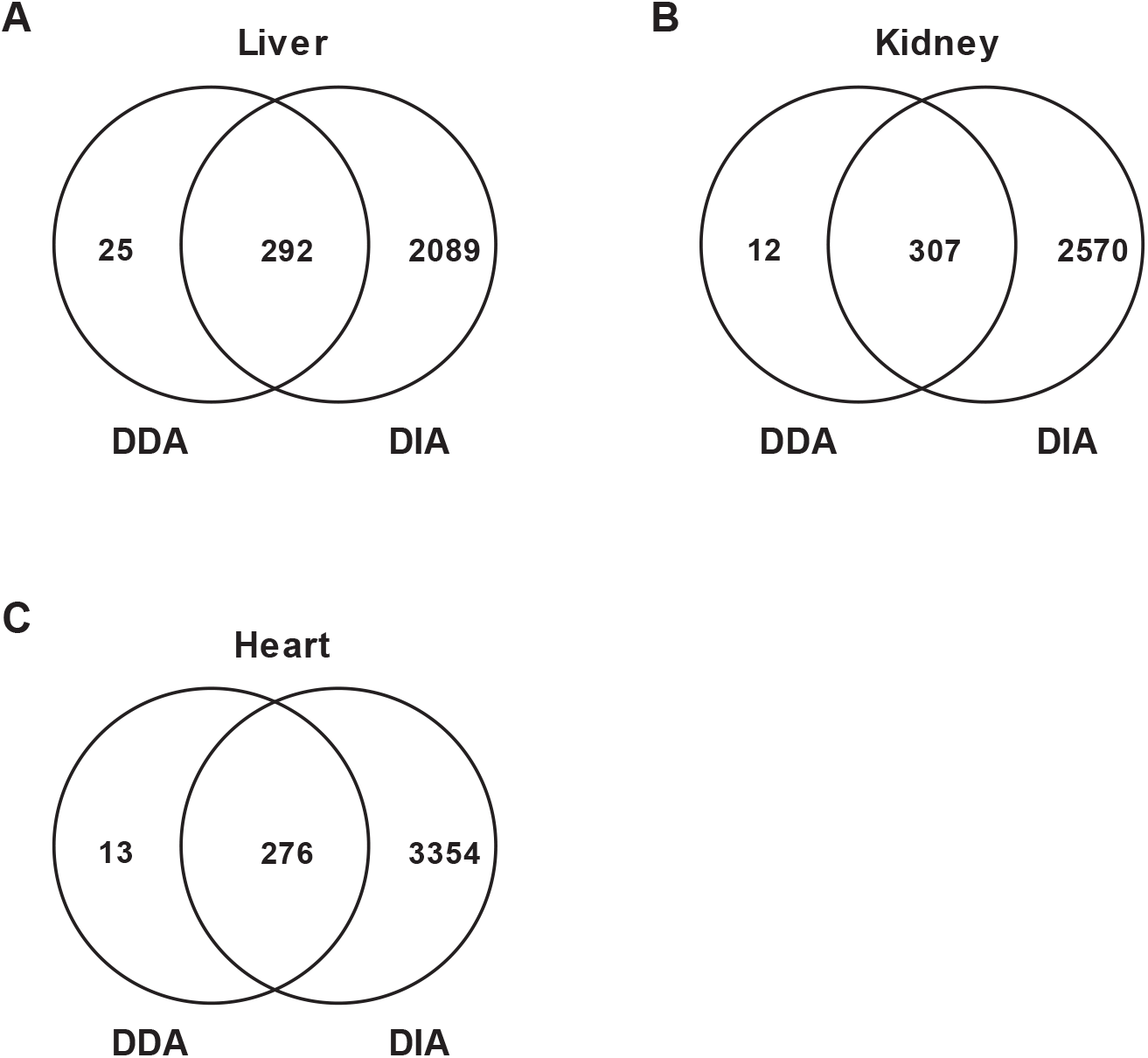
Comparison of the vascular cell surface proteins previously identified using a standard shotgun DDA-proteomics workflow (17) and in this study using a DIA-SWATH-MS pipeline. Venn diagrams showing unique and shared protein identifications in (A) liver, (B) kidney and (C) heart.

**Fig. S3.**
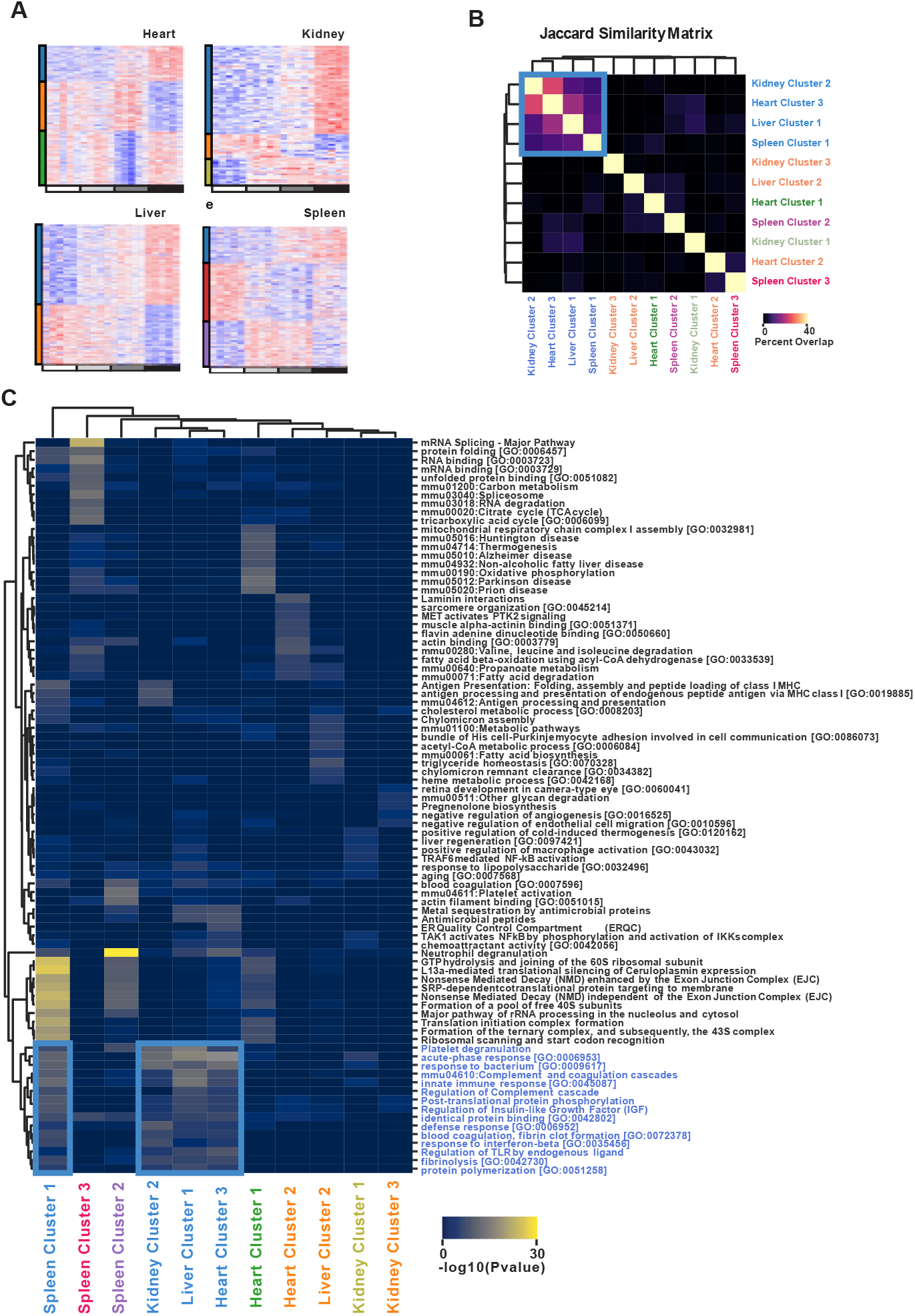
Coordinated parenchymal proteome responses triggered by S. aureus sepsis. *(A)* Heatmaps showing the k-means clustering of significant (1-way ANOVA, FDR <0.10) parenchymal protein identifications in the heart (n=5), kidney (n=5), spleen (n=5), and liver (n=5). *(B)* Jaccard similarity matrix (intersection over the union) representing the overlap of the protein identifications across parenchymal clusters derived from the proteomics analysis. *(C)* Functional enrichment analysis of the parenchymal proteome clusters. Hypergeometric test for functional enrichment was used and colored according to the P-value. Hierarchical clustering was utilized to visualize functional similarities between the indicated protein clusters. Clusters with the same proteome trajectory (i.e., temporal intensity patterns) are coded with the same color as represented in the bars to the left of the heatmaps.

## Supplementary dataset legends

**Supplementary Dataset. 1.** Normalized protein intensities for enriched vascular proteomes. The whole quantified proteome for the enriched liver, kidney, heart, and spleen vasculature.

**Supplementary Dataset. 2**. Normalized protein intensities for plasma and parenchymal proteomes. The whole quantified proteome for plasma, liver, kidney, heart, and spleen. Both intensities harvested from the DDA and DIA approaches are listed for comparison.

## References

1. M. Singer, et al., The third international consensus definitions for sepsis and septic shock (sepsis-3). JAMA - J. Am. Med. Assoc. 315, 801–810 (2016).

2. K. E. Rudd, et al., Global, regional, and national sepsis incidence and mortality, 1990– 2017: analysis for the Global Burden of Disease Study. Lancet 395, 200–211 (2020).

3. J. C. Marshall, Why have clinical trials in sepsis failed? Trends Mol. Med. 20, 195–203 (2014).

4. C. Pierrakos, D. Velissaris, M. Bisdorff, J. C. Marshall, J. L. Vincent, Biomarkers of sepsis: Time for a reappraisal. Crit. Care 24, 1–15 (2020).

5. C. Lelubre, J. L. Vincent, Mechanisms and treatment of organ failure in sepsis. Nat. Rev. Nephrol. 14, 417–427 (2018).

6. A. Leligdowicz, M. Richard-Greenblatt, J. Wright, V. M. Crowley, K. C. Kain, Endothelial activation: The Ang/Tie axis in sepsis. Front. Immunol. 9, 1–15 (2018).

7. S. Han, et al., Amelioration of sepsis by TIE2 activation-induced vascular protection. Sci. Transl. Med. 8, 1–12 (2016).

8. J. Hauschildt, et al., Dual Pharmacological Inhibition of Angiopoietin-2 and VEGF-A in Murine Experimental Sepsis. J. Vasc. Res. 57, 34–45 (2020).

9. J. D. Lapek, et al., Defining Host Responses during Systemic Bacterial Infection through Construction of a Murine Organ Proteome Atlas. Cell Syst. 6, 579-592.e4 (2018).

10. N. H. Tran, et al., Complete De Novo Assembly of Monoclonal Antibody Sequences. Sci. Rep. 6, 31730 (2016).

11. E. Malmström, et al., Large-scale inference of protein tissue origin in gram-positive sepsis plasma using quantitative targeted proteomics. Nat. Commun. 7 (2016).

12. S. Michalik, et al., Causes Changes in the Concentrations of Lipoproteins and Acute-Phase Proteins and Is Associated with Low Antibody Titers against Bacterial Virulence Factors. 5, 1–17 (2020).

13. J. M. Wozniak, et al., Mortality Risk Profiling of Staphylococcus aureus Bacteremia by Multi-omic Serum Analysis Reveals Early Predictive and Pathogenic Signatures. Cell 182, 1311-1327.e14 (2020).

14. G. Pimienta, et al., Plasma Proteome Signature of Sepsis: a Functionally Connected Protein Network. Proteomics 19 (2019).

15. A. C. A. Cleuren, et al., The in vivo endothelial cell translatome is highly heterogeneous across vascular beds. Proc. Natl. Acad. Sci. U. S. A. 116, 23618–23624 (2019).

16. E. Durr, et al., Direct proteomic mapping of the lung microvascular endothelial cell surface in vivo and in cell culture. Nat. Biotechnol. 22, 985–992 (2004).

17. A. G. Toledo, et al., Proteomic atlas of organ vasculopathies triggered by Staphylococcus aureus sepsis. Nat. Commun. 10, 4656 (2019).

18. G. J. Golden, et al., Endothelial Heparan Sulfate Mediates Hepatic Neutrophil Traf fi cking and Injury during Staphylococcus aureus Sepsis.

19. R. Bruderer, et al., Optimization of experimental parameters in data-independent mass spectrometry significantly increases depth and reproducibility of results. Mol. Cell. Proteomics 16, 2296–2309 (2017).

20. V. Demichev, C. B. Messner, S. I. Vernardis, K. S. Lilley, M. Ralser, DIA-NN: neural networks and interference correction enable deep proteome coverage in high throughput. Nat. Methods 17, 41–44 (2020).

21. L. Daniel D. S. H. Sebastian, Learning the parts of objects by non-negative matrix factorization. Nature 401, 788–791 (1999).

22. C. von Mering, et al., STRING: A database of predicted functional associations between proteins. Nucleic Acids Res. 31, 258–261 (2003).

23. H. Tang, et al., Sepsis-induced coagulation in the baboon lung is associated with decreased tissue factor pathway inhibitor. Am. J. Pathol. 171, 1066–1077 (2007).

24. J. Taylor, K. Reinhart, B. Vallet, Staging of the pathophysiologic responses of the primate microvasculature to Escherichia coli and endotoxin: Examination of the elements of the compensated response and their links to the corresponding uncompensated lethal variants. Crit. Care Med. 29 (2001).

25. P. F. Laterre, et al., A clinical evaluation committee assessment of recombinant human tissue factor pathway inhibitor (tifacogin) in patients with severe community-acquired pneumonia. Crit. Care 13 (2009).

26. E. Abraham, et al., Efficacy and Safety of Tifacogin (Recombinant Tissue Factor). JAMA 290, 238–247 (2003).

27. Wang Y, et al. Modeling human congenital disorder of glycosylation type IIa in the mouse: Conservation of asparagine-linked glycan-dependent functions in mammalian physiology and insights into disease pathogenesis. Glycobiology.11:1051–1070. 2001)

28. F. Desiere, et al., The PeptideAtlas project. Nucleic Acids Res. 34, 655–658 (2006).

29. C. Q. Zhong, et al., Generation of a murine SWATH-MS spectral library to quantify more than 11,000 proteins. Sci. Data 7, 1–9 (2020).

30. D. W. Huang, B. T. Sherman, R. A. Lempicki, Systematic and integrative analysis of large gene lists using DAVID bioinformatics resources. Nat. Protoc. 4, 44–57 (2009).

31. F. Supek, M. Bošnjak, N. Škunca, T. Šmuc, Revigo summarizes and visualizes long lists of gene ontology terms. PLoS One 6 (2011).

32. A. L. Tarca, V. J. Carey, X. wen Chen, R. Romero, S. Drǎghici, Machine learning and its applications to biology. PLoS Comput. Biol. 3 (2007).

33. V. D. Blondel, J. L. Guillaume, R. Lambiotte, E. Lefebvre, Fast unfolding of communities in large networks. J. Stat. Mech. Theory Exp. 2008 (2008).

34. Paul Shannon, et al., Cytoscape: A Software Environment for Integrated Models. Genome Res. 13, 426 (1971).

